# Ly6C^hi^ monocytes are metabolically reprogrammed in the blood during inflammatory stimulation allowing for macrophage lineage commitment

**DOI:** 10.1101/2021.11.15.468584

**Authors:** Gareth S.D. Purvis, Eileen McNeill, Benjamin Wright, Santiago Revale, Helen Lockstone, Keith M. Channon, David R. Greaves

**Affiliations:** Sir William Dunn School of Pathology, University of Oxford, Oxford, UK; Wellcome Trust Centre for Human Genetics, University of Oxford, Oxford, UK; Division of Cardiovascular Medicine, Radcliffe Department of Medicine, University of Oxford, Oxford, UK; British Heart Foundation Centre of Research Excellence, University of Oxford, Oxford, UK; NIHR Oxford Biomedical Research Centre, Oxford, University Hospitals NHS Foundation Trust, John Radcliffe Hospital, Oxford, OX3 9DU

## Abstract

Acute inflammation is a rapid and dynamic process involving the recruitment and activation of multiple cell types in a co-ordinated and precise manner. Using cell tracking, linage tracing and single cell transcriptomics we investigated the origin and transcriptional reprogramming of monocytes and macrophages in acute inflammation.

Monocyte trafficking and adoptive transfer experiments revealed that monocytes undergo rapid phenotypic change as they exit the blood and give rise to monocyte-derived macrophages that persist during the resolution of inflammation. Single cell transcriptomics revealed significant heterogeneity within the surface marker defined CD11b^+^Ly6G^-^Ly6C^hi^ monocyte population within the blood and at the site of inflammation. Lineage trajectory analysis revealed that Ly6C^hi^ monocytes in the blood are re-programmed into a defined differentiation pathway following inflammatory stimulus. We show that two major transcriptional reprogramming events occur during the initial 6 h of Ly6C^hi^ monocyte mobilisation, one in the blood priming monocytes for migration and a second at the site of inflammation. Pathway analysis revealed an important role for oxidative phosphorylation (OxPhos) during both these reprogramming events in a subset of M2-like cells. Experimentally we also demonstrate that OxPhos is essential for murine and human monocyte chemotaxis. These new findings opening up the possibility that altering monocyte metabolic capacity towards OxPhos could facilitate enhanced macrophage M2-like polarisation to aid inflammation resolution and tissue repair.

## Introduction

As an acute inflammatory response evolves, the cellular composition at the site of inflammation rapidly changes. The initial recruitment of polymorphonuclear leukocytes (PMN’s) is overtaken by in an influx of monocytes from the circulation. Classically monocytes are divided as three main classes in humans and mice: classical (CD14^+^CD16^−^ in humans and Ly6C^hi^ in mice), intermediate (CD14^+^CD16^+^ in humans and Ly6C^+^Treml4^+^ in mice), and nonclassical (CD14^−^CD16^+^ in humans and Ly6C^lo^ in mice) (Babu Narasimhan et al., 2019). During the inflammatory response monocytes are released into the blood from two main stores, the bone marrow which is the origin of all haemopoietic lineages, and from the spleen (Guilliams et al., 2018)(Swirski et al., 2009). This process is tightly regulated through the release of specific chemoattractants and the expression of their cognate receptors on target cells (White et al., 2013).

Once released from either bone marrow or splenic chemokine:chemokine receptor interactions lead blood monocytes to the site of inflammation. To date little is known about the metabolic demand for monocyte chemotaxis (Marelli-Berg and Jangani, 2018). Following peritoneal challenge with zymosan, in the mouse, intermediate Ly6C^+^Treml4^+^ and classical Ly6C^hi^ monocytes are rapidly recruited to the inflamed peritoneum. Our current understanding suggests that monocytes utilise glycolytic metabolism after activation, however, most studies measure this *ex vivo* and using lipopolysaccharide as the stimulus (Lee et al., 2019)(Kishore et al., 2017). Non-classical Ly6C^lo^ monocytes which tend to have a patrolling function and show less recruitment to the peritoneum (Merah-Mourah et al., 2020)(Iqbal et al., 2016); little is known about their energy utilisation. Monocytes that are recruited to a site of inflammation can undergo differentiation into dendritic cells or macrophages.

In the later stages of the inflammatory response macrophages are the predominant cell type present. However, the exact origin of ‘resolution phase’ macrophage populations has been difficult to pin down and their function are less studied (Watanabe et al., 2019). The classical paradigm of M1 (pro-inflammatory) and M2 (pro-resolution) macrophage differentiation is currently being challenged, with limited evidence of distinct classes of macrophages described *in vitro* being present *in vivo* (Nahrendorf and Swirski, 2016). It is likely that macrophage differentiation *in vivo* is more dynamic and complex than macrophage differentiation *in vitro* (Martinez and Gordon, 2014). Tissue resident macrophage populations are classically thought of as ‘M2-like’ as they harbour homoeostatic functions (Murray et al., 2014). The peritoneal cavity contains two populations of resident macrophages. The predominant cell type is large peritoneal macrophage (LPM) that are embryonically seeded and long-lived. There are also a more rare population of short-lived MHCII^+^ monocyte-derived macrophage termed small peritoneal macrophages (SPM) (Ghosn et al., 2010)(Bain et al., 2016).

Monocyte-derived macrophages recruited following inflammatory stimulus exhibit distinct transcriptional, functional and phenotypic signatures to resident cells. In the peritoneal cavity, sterile inflammation can cause substantial loss in number of LPM through egress, cell death or loss in fibrin clots (Gautier et al., 2013) but the extent of this loss appears dependent on stimulus and severity of the ensuing inflammation (Davies et al., 2013); however, the LPM can be replenished through proliferation during the resolution phase (Davies et al., 2011). During peritoneal inflammation including that caused by abdominal surgery there can be loss of resident cells, which are partially replaced by bone marrow derived cells; again with the degree of replacement correlates to the extent of initial loss (Bain et al., 2020)(Louwe et al., 2021).

Current models suggest recruited blood monocytes differentiate into inflammatory M1-like macrophages upon entering the site of inflammation following activation from local stimuli i.e. bacterial cell wall component or oxLDL in the atherosclerotic lesion (Auffray et al., 2009); and that over time the M1-like macrophages in resolving inflammation transition to become more M2-like. Recent work demonstrated M(LPS/IFNγ) bone marrow-derived macrophage (BMDM) cannot undergo phenotypic switch to become M2-like unless iNOS is inhibited (Van Den Bossche et al., 2016). Inhibition of NO production resulted in M(LPS/IFNγ) macrophages not undergoing the Warburg metabolic switch (Bailey et al., 2019); having an intact electron transport chain meant upon IL-4 stimulation macrophages could undergo M2 differentiation (Van Den Bossche et al., 2016). These data suggest macrophages have limited ability to undergo an M1 to M2 phenotypic switch *in vivo*. One key feature consistent with all models is that macrophage metabolic state is a key indicator of their activation state (Viola et al., 2019).

In this study we sought to investigate if there is heterogeneity within circulating and recruited Ly6C^hi^ monocyte populations, and to identify the earliest pathways that regulate monocyte recruitment and differentiation at sites of zymosan-induced peritonitis. Single cell transcriptomics allowed us to compare the transcriptome of monocytes in the blood (naïve to inflammatory stimuli), with newly recruited monocytes (in the peritoneum post inflammatory stimuli) without the need for tissue digestion and lengthy sorting protocols. This experimental design provided unique insights in to how circulating blood Ly6C^hi^ monocytes are reprogrammed during acute resolving inflammation in a spaciotemporal manner at a single cell resolution. Our study revealed 1) there is previously underappreciated heterogeneity within Ly6C^hi^ blood; 2) that Ly6C^hi^ monocyte cellular fate at a site of inflammation is pre-determined in the blood 3) a sub-set of Ly6C^hi^ monocytes which are pre-programmed to become M2-like macrophages are dependent on a metabolic switch towards oxidative phosphorylation as early as 2 h post inflammatory stimulation in the blood and 4) OxPhos via intact mitochondrial electron transport chain is critical for monocytes chemotaxis.

## Methods

### Animal ethics

All animal studies were conducted with ethical approval from the Local Ethical Review Committee at the University of Oxford and in accordance with the UK Home Office regulations (Guidance on the Operation of Animals, Scientific Procedures Act, 1986). Mice were housed in ventilated cages with a 12-h light/dark cycle and controlled temperature (20–22 °C), and fed normal chow and water ad libitum. For in vivo experiments adult mice >10 weeks of age were used for all studies. Mice had received no prior procedures (including acting as breeding stock) prior to use in experiments for this manuscript.

### Zymosan induced peritonitis

Mice were injected with 100 μg zymosan A (Sigma) in 500 μl PBS using an 30G insulin syringe. At defined time points mice underwent peritoneal lavage post-mortem with ice-cold PBS/EDTA (5 mM), lavage fluid was injected using a 25 G needle, the cavity mixed by agitation, and then lavage fluid was withdrawn using a 21 G needle. Cellular recruitment was assessed by flow cytometry using an absolute cell count method as previously described (Purvis et al., 2021). Total cellular recruitment was calculated as the total cells that would be recovered in the 5 ml lavage volume.

### Adoptive transfer

Monocytes were isolated from *CD68*-GFP bone marrow as described previously (McNeill et al., 2017). Briefly, bone marrow cell suspensions were passed through a 70 μm cell strainer and red blood cells lysed (LCK Buffer, Sigma) for 10 mins on ice. The bone marrow monocytes were isolated using the Murine Bone Marrow Monocyte Isolation kit (Miltenyi Biotec) and underwent negative selection using an autoMACS Pro Separator (Miltenyi Biotec). The purity of the resulting populations confirmed by flow cytometry. Bone marrow isolations from a total of 2 femurs would typically yield 2 × 10^6^ cells at a purity of 90% monocytes.

Isolated h*CD68*-GFP monocytes (1 × 10^6^, in 200 μl) were delivered intravenously 30 mins after C57BL/6J mice were administered 100 μg zymosan A in PBS (Sigma) *i*.*p*. After 24 or 48 hr mice were sacrificed and peritoneal exudates were collected by peritoneal lavage in ice-cold PBS/EDTA (5 mM). Total cell counts and cellular composition of peritoneal exudate were determined by flow cytometry full gating strategy found in Supplementary figure 1.

### Cell capture, cDNA synthesis, single cell RNA-Seq library preparation

Whole blood and peritoneal exudate cells were isolated from mice. Red blood cells lysed (LCK Buffer, Sigma) for 10 mins on ice and cells were stained for flow cytometry. Briefly, cells were live/dead stained (IR Live Dead Stain, Thermofischer) was added and washed after 10 mins. Cells were incubated with Fc-blocking anti-CD16/32 anti-bodies followed by cell hashing antibody (1 μg/ml, BioLegend Cell Hashing A) (Stoeckius et al., 2018) combined and stained with anti-CD11b-PerCP, Ly6G-FITC and Ly6C-PE antibodies (BioLegend, USA). The cell hashing approach allowed each of 5 biological replicates from the 4 experimental groups to be combined and sorted in one sample per experimental group, whilst still capturing data from each individual mouse.

Stained cells were then sorted and CD11b^+^LygG^-^Ly6C^+^ monocytes were collected and washed in PBS with 0.04 % BSA and re-suspended at a concentration of ∼50 cells/μl before capturing single cells in droplets on Chromium 10x Genomics platform. Library generation for 10x Genomics v3.0 chemistry was performed following the Chromium Single Cell 3′ Reagents Kits User Guide: v3.0 rev A, CG000183 (10X Genomics). Quantification of cDNA was performed using Qubit dsDNA HS Assay Kit (Life Technologies, Q32851) and Agilent High Sensitivity D5000 ScreenTape (Agilent, 5067-5592). Quantification of library was performed using Qubit dsDNA HS Assay Kit (Life Technologies, Q32851) and D1000 ScreenTape (Agilent. 5067-5582). Libraries were prepared and sequenced, with an average of 38628±5029 reads per cells.

### Sequencing and Base Calling

Both the 10X Single Cell RNAseq 3’ v3 and the Cell Hashing libraries were sequenced together on a single Illumina HiSeq 4000 lane at the Oxford Genomics Centre using a 28×98bp paired-end read configuration. Base-calling and demultiplexing were performed using “cellranger mkfastq” pipeline from the 10X Genomics Cell Ranger Suite (v3.0.2) (10X Genomics) in order to produce Cell Ranger compatible FastQ files.

### Quality Control and Processing

FastQ files were quality controlled using FastQC (Andrews et al., 2010). Gene expression and cell hashing libraries originating from the same sample were processed together using “cellranger count” pipeline with the “--libraries” option and default arguments. Data were aligned against the pre-built mouse reference available from 10X Genomics (10X Genomics) which uses GRCm38 (Ensembl 93) primary assembly and annotation files. Cell Ranger’s “count” pipeline performs the alignment, filtering, barcode counting, and UMI counting processes. It uses the Chromium cellular barcodes to generate a feature-barcode matrix, and because both the gene expression and cell hashing libraries were provided together, the matrix includes both the gene expression and cell hashing feature counts. Cell hashing efficiency was assessed with ridge plots analysis and 5 distinct signatures detected allowing for biological variation within experimental group to be factored into all subsequent analysis (Sup Figure 2).

### Single-Cell Analyses

The previously generated gene expression and cell hashing counts matrix was parsed using Seurat v3 (Stuart et al., 2019). Following exclusion of doublets and cells with aberrant ribosomal RNA to genomic RNA 8389 cells were included in the subsequent analysis. Cells were assigned to clusters using Seurat and the expression profiles for the clusters were examined to determine if any were cells of an unexpected type. Such cells were excluded from the data set and the clusters recalculated. The reclustered data was taken forward for subsequent analysis. Characteristic markers for the new clusters were established also using Seurat.

### Pseudotime Analysis

The Monocle3 R package (Trapnell et al., 2014)(Qiu et al., 2017a)(Qiu et al., 2017b)(Cao et al., 2019) was used to estimate a pseudotime progression for the cells. Initial analysis produced disjoint pseudotime progressions across the different clusters found. The data set was then subdivided iteratively until each pseudotime model showed one progression for all cells in that set. Each pseudotime model requires the specification of a set of ‘root’ cells assumed to be the start of the pseudotime progression. This was taken to be those cells in the model belonging to the naive blood group, if any were present. If not, then the Blood 2hr cells. If no cells belonging to either group were present, PEC 2hr was taken as the set of root cells (this was for pseudotime models I-VIII). Alternative models used the cluster analysis to establish the set of root cells for further refinement (models IX-XI).

### Generation of murine bone marrow-derived macrophages (BMDM)

Bone marrow was obtained by flushing the femur and tibia of adult female mice with PBS. A single cell suspension was prepared by passing the bone marrow through a 70 μm cell strainer. Whole bone marrow cultures were then cultured in 10 cm non-tissue culture treated dishes for 7 days in Dulbecco’s modified Eagle’s medium (DMEM) containing 25 mM glucose (Invitrogen) and supplemented with 100 U/ml penicillin and 100 ng/ml streptomycin (Sigma), 10 % (v/v) fetal bovine serum (PAA Laboratories) and 10 % (v/v) L929 (ATCC NCTC clone 929) conditioned medium at 37 °C and 5 % CO_2_. Monocyte to macrophage differentiation experiments, bone marrow monocytes were immuno-magnetically isolated and culture for 7 days in Dulbecco’s modified Eagle’s medium (DMEM) containing 25 mM glucose (Invitrogen) and supplemented with 100 U/ml penicillin and 100 ng/ml streptomycin (Sigma), 10 % (v/v) fetal bovine serum (PAA Laboratories) and 10 % (v/v) L929 (ATCC NCTC clone 929) conditioned medium at 37 °C and 5 % CO_2._ The protocol for macrophage culture was validated by flow cytometry analysis of the differentiated cells using F4/80 and CD11b as macrophage cell markers (Purvis et al., 2020).

### Isolation and preparation of human monocyte derived monocytes (hMoDM)

Peripheral blood mononuclear cells (PBMC) were purified from leukocyte cones from healthy volunteers with informed consent (NHSBTS, Oxford, UK) by density centrifugation over Ficoll-Paque PLUS (Sigma). The PBMC layer was carefully harvested and then washed twice with PBS. Monocytes were then isolated from the PBMCs by negative selection using magnetic beads (Miltenyi Biotec, Bergisch Gladbach, Germany). Cells were maintained in RPMI 1640 medium supplemented with 1 % human serum, 50 ng/ml macrophage colony stimulating factor (hM-CSF, BioLegend, San Diego, USA), 100 U/ml penicillin and 100 ng/ml streptomycin (Sigma) for 7 days.

### Flow cytometry

Cells were washed in FACS buffer (0.05 % BSA, 2 mM EDTA in PBS pH 7.4) blocked using anti CD16/32 for 10 mins at 4°C, followed by antibody staining for the surface markers. The gating strategy for assessing myeloid cell composition of peritoneal exudates is found in Sup Figure 1. Absolute cell counts were performed using a defined quantity of calibration beads added to each sample (CountBright, Invitrogen). Data were acquired using either a DAKO CyAn cytometer and Summit software (both Beckton Coulter) or a BD Fortessa X20 cytometer and Diva software (both BD Biosciences) and then analysed using FlowJo (Tree Star Inc, USA) software.

### ACEA xCELLigence real-time cell migration

Experiments were carried out with CIM-16 plates and an xCELLigence RTCA-DP instrument (ACEA, San Diego, USA) as previously described (Iqbal et al., 2016). Briefly, chemoattractants were made to desired concentrations (C5a, 10 nM, CCL2 10 nM) and loaded into the lower wells of the CIM-16 plate. Upper wells were filled with chemotaxis buffer and plates equilibrated for 30 min at RT. Freshly isolated murine bone marrow monocytes, human monocytes or BMDM’s were resuspended in chemotaxis buffer and incubated with OxPhos inhibitors 30 min at 37°C, 5% CO_2_. Cell suspensions were placed into the wells of the upper chamber, and the assay performed over 6 hr (readings were taken every 15 s).

## Results section

### Single cell transcriptomics reveals heterogeneity within Ly6C^hi^ monocyte populations

Following a 100 μg zymosan challenge Ly6C^hi^ monocytes are recruited to the peritoneum (Figure 1A) and within 48 h a new population of macrophages appear (CD11b^+^Ly6C^+^CD115^-^ to CD11b^+^Ly6C^-^CD115^+^F4/80^lo^) (Figure 1B). To confirm monocytes are recruited from the circulation we next performed intravenous monocyte adoptive transfer of h*CD68*-GFP expressing bone marrow monocytes (Iqbal et al., 2014) into ongoing zymosan induced peritonitis (Figure 1C) and demonstrated that circulating h*CD68*-GFP monocytes were recruited to the peritoneum from the circulation (Sup Figure 3D) and differentiate into monocyte-derived macrophages in line with host cell at the site of inflammation within 48 h (Figure 1E). These experiments confirm that the peritoneal cavity is repopulated with monocyte-derived macrophages, following zymosan challenge.

**Figure 1:**
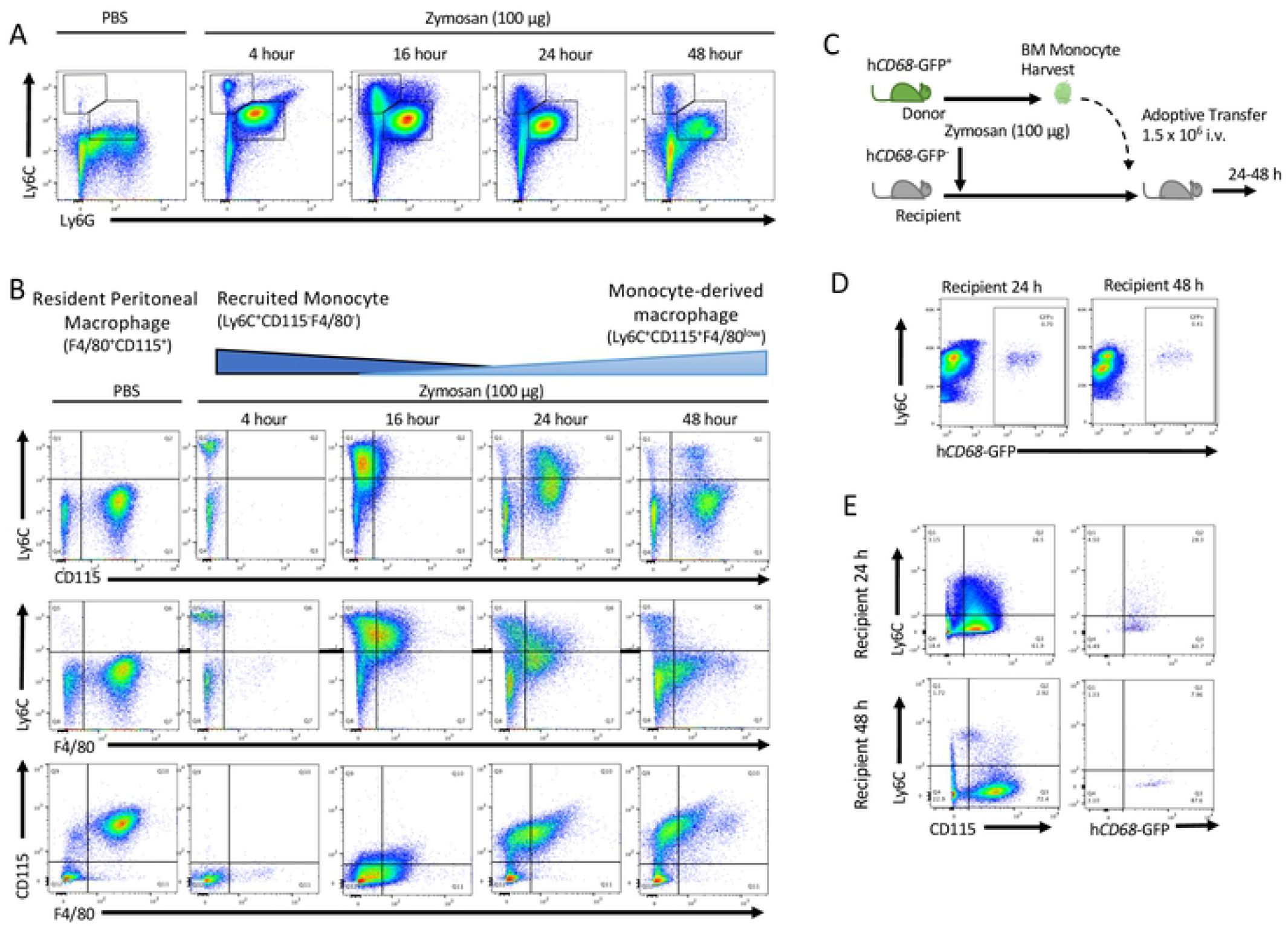
Resolution phase macrophages differentiate from blood derived monocytes in zymosan induced peritonitis. hCD68-GFP monocytes are recruited to the peritoneum and differentiate into monocyte derived macrophages following zymosan challenge. Peritoneal exudate were harvested from h*CD68*-GFP 4 h, 16, 24 and 48 h after zymosan challenge (100 ug; *i*.*p*.) and cellular composition analysed by flow cytometry. **(A)** Representative flow cytometry of peritoneal exudate monocytes (Ly6G^-^Ly6C^+^) and neutrophils (Ly6G^+^Ly6C^+^) which have been first gated on h*CD68*-GFP^+^CD11b ^+^ cells. **(B)** Schematic showing the temporal changes in myeloid cell surface markers between resident peritoneal macrophage, recruited monocyte and monocyte-derived macrophage. Representative flow cytometry of monocyte to macrophages differentiation following zymosan challenge (100 ug; *i*.*p*.) using three separate sets of cells surface antigens Ly6C vs. CD115, Ly6C vs. F4/80 and CD115 vs. F4/80 after being gated on h*CD68*-GFP^+^ cells. **(C)** Schematic showing adoptive transfer of bone marrow monocytes from donor hCD68-gfp^+^ mice into recipient hCD68-gfp^-^ following zymosan challenge (100 ug; *i*.*p*.). **(D)** Representative flow cytometry of peritoneal exudates 24 h and 48 h after adoptive transfer of h*CD68*-GFP^+^ bone marrow monocytes showing monocyte to macrophages differentiation and h*CD68*-GFP expression. Representative data of n=3 independent experiments.

To capture the earliest transcriptional reprogramming of circulating and recruited Ly6C^hi^ monocytes we used a single cells RNA sequencing approach. We chose 3 timepoints and isolated either circulating or peritoneally recruited monocytes: 0 h (Naïve blood); 2 h (Blood and PEC 2h) and 6 h (PEC 6h) after zymosan challenge, as at these time points there was monocyte recruitment to the peritoneum, but it was prior to alterations in their cell surface markers. This allowed fluorescent antigen cell sorting (FACS) to isolate the same population of CD45^+^CD11b^+^Ly6G^-^Ly6C^hi^ monocytes (Figure 2A). Following single cell capture and RNA-sequencing principal component analysis revealed the transcriptomes from biological replicates segregated together in clusters based on experimental group (Figure 2B).

**Figure 2:**
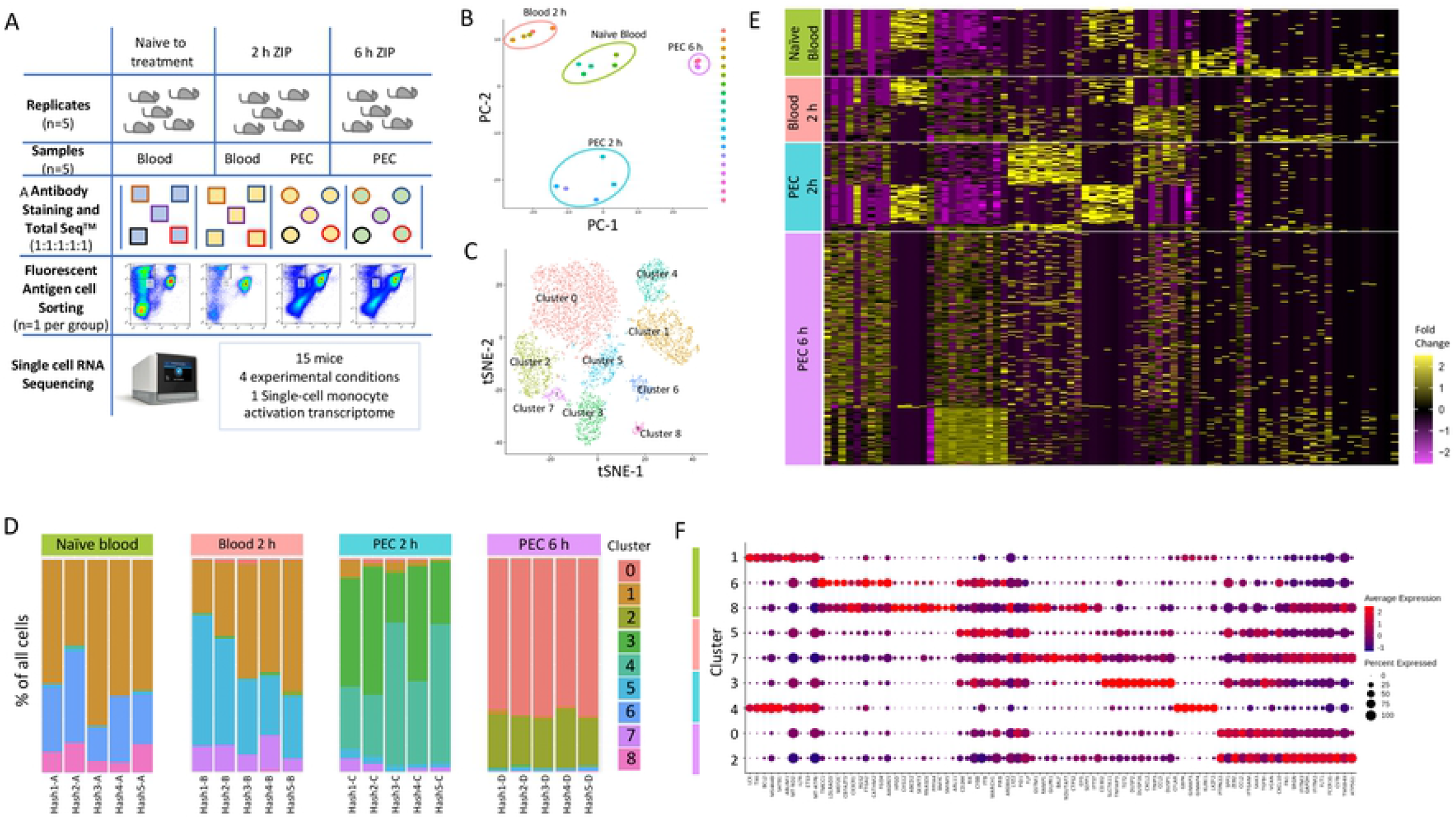
Single cell RNA-Sequencing to profile Ly6C^hi^ monocytes following zymosan induced peritonitis reveals significant heterogeneity. **(A)** Workflow and experimental scheme including experimental conditions (naïve to treatment, 2 h zymosan induced peritonitis (ZIP) and 6 h ZIP), cell types harvested (blood or Peritoneal exudates), cells sorted (CD11b^+^Ly6G^-^Ly6C^+^) and processing workflow (Cell hashing and 10X Genomics followed by Illumina High Seq 4000). **(B)** PCA plot of bulk scRNA-seq profiles for each sample. **(C)** *t*-SNE visualization of 8393 cells following exclusion criteria, coloured by cluster identity from Louvain clustering. **(D)** Bar chart showing the percentage of cells per cluster identified based on cell hashing identity. **(E)** Heatmap of differential gene expression of cells vs all other cells, and grouped based on experimental group (Naïve blood (green), 2 h blood (salmon), 2 h PEC (turquoise) and 6 h PEC (lilac). **(F)** Dot plot showing expression of top 10 genes per cluster.

We subjected 5,825 single cell transcriptomes to Louvain clustering and *t*-SNE visualisation (Figure 2C). Within the 4 experimental groups, Naïve blood, Blood 2h, PEC 2 h and PEC 6 h there are 9 distinct clusters. Across all 9 clusters 11,880 genes were significantly upregulated and 6403 genes significantly down regulated more than 2-fold compared to each other cluster (5 % false discovery rate) (Sup Figure 3A).

We observed 3 distinct clusters within the naïve blood monocytes (Figure 2D). Within the blood 2 h monocytes there were also three clusters, 2 new clusters and cells which fell into cluster 1 which is shared with naïve blood (Figure 2D). Within the PEC 2 h monocytes there was the emergence of two further distinct clusters (Figure 2D). And within the PEC 6 h monocytes there were two further unique clusters (Figure 2D). Remarkably, there was only limited biological variation between mice within each experimental group, but significant biological variation between groups (Figure 2D). Our experiments revealed previously unknown heterogeneity within the CD11b^+^Ly6G^-^Ly6C^hi^ monocyte population. Differential gene expression revealed a pattern of rapid and divergent differentiation in the blood Ly6C^hi^ monocytes as early as two hours after zymosan challenge, but also showed that Ly6C^hi^ monocytes recruited to the peritoneum continued on a differentiation pathway characterised by differential gene expression of discrete subsets of genes (Figure 2E). Each cluster had a unique transcriptional profile, but as all the cells analysed are Ly6C^hi^ monocytes it was not unsurprising that each cluster did not have unique gene identifiers, indeed there was significant overlap in markers between clusters (Figure 2F).

### Ly6C^hi^ monocyte recruitment and differentiation to monocyte-derived macrophages *in vivo* leads to the activation of multiple transcriptional pathways

Having demonstrated heterogeneity within the Ly6C^hi^ monocyte population we next wanted to understand the pathways which were differentially regulated as monocytes are recruited from the blood to the peritoneum within the first 6 h following inflammatory insult. Pseudo-bulk analysis of monocyte clusters demonstrated that Ly6C^hi^ monocytes segregated by experimental group (time point) (Figure 3A). This was indicative of divergent transcriptomes during early activation. We have fully characterised for the first time, at a single cell level the temporal gene expression profiles of Ly6C^hi^ monocytes in the first 6-hours after zymosan challenge *in vivo* (Figure 3A). We identified biological processes associated with monocyte re-programming within the blood and the peritoneum having performed pathway enrichment analysis using Ingenunity Pathways Analysis (IPA). Two hours after zymosan stimulation key pathways that were are up-regulated in 2 h blood monocytes compared to naïve blood monocytes with a z-score greater than 2.5 included those related to monocyte and macrophage acute response including NO production, FcγR mediated phagocytosis and pathogen pattern recognition receptors. Pathways that were also enriched included those needed for cellular motility including RhoA and Cdc42 signalling. The top-ranked enriched canonical pathway was oxidative phosphorylation (OxPhos) (Figure 3B).

**Figure 3:**
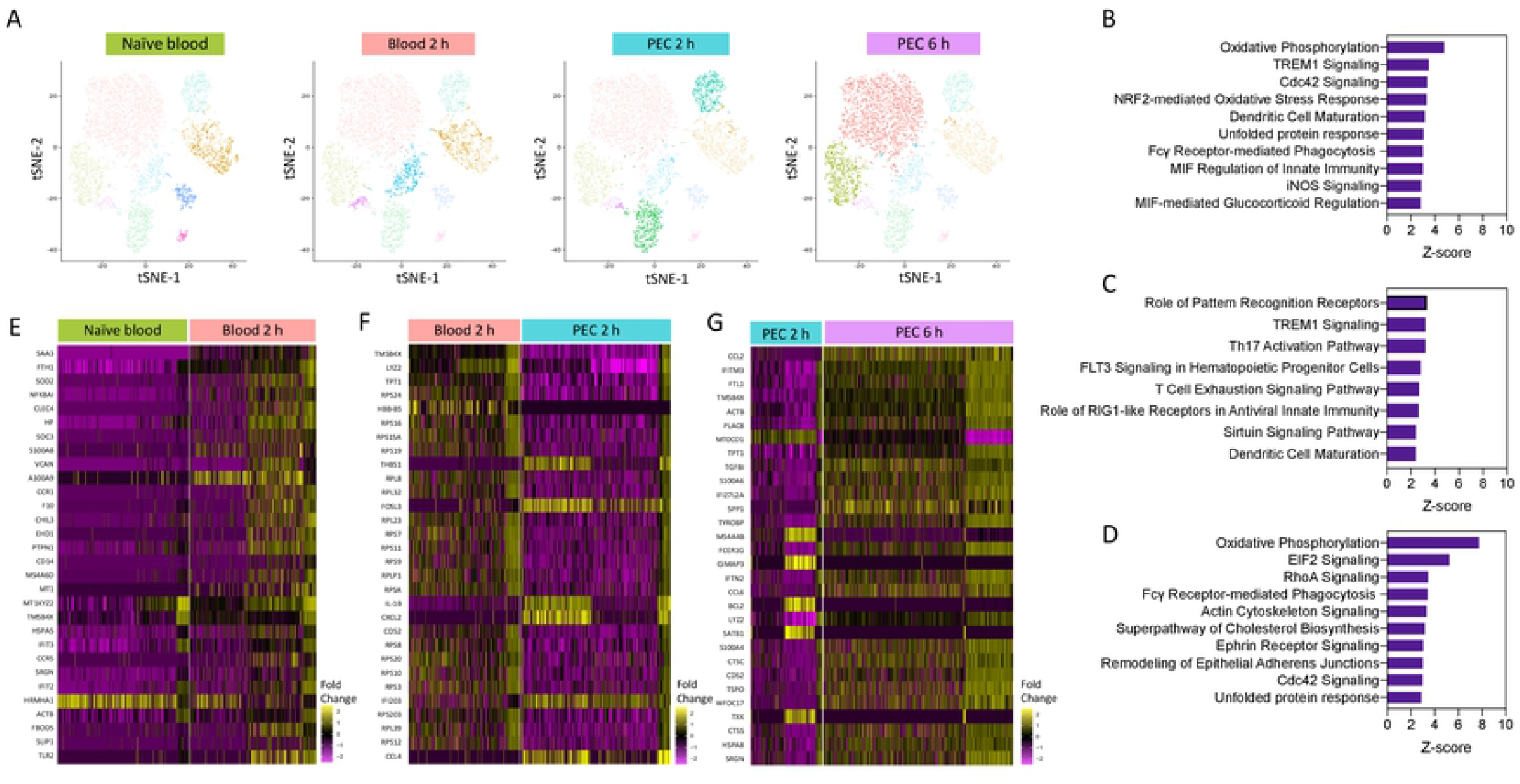
Pathways enrichment analysis reveals a central role for oxidative phosphorylation. Pathway enrichment analysis reveals a central role for oxidative phosphorylation in Ly6C^hi^ monocyte activation. **(A)** tSNE visualisation of cells based on cluster, and highlighted cells present in each experimental condition. Pathway enrichment analysis between **(B)** 2h blood and naïve blood, **(C)** 2 h PEC and 2 h blood and **(D)** 6 h PEC and 2 h PEC. Heap map of the top 30 differentially expressed genes between **(E)** 2h blood and naïve blood, **(F)** 2 h PEC and 2 h blood and **(G)** 6 h PEC and 2 h PEC.

Pathway analysis revealed that when compared to 2 h blood monocytes, 2 h PEC monocytes significantly up regulated pathways with a Z-score less than 2.5 were associated with Pattern Recognition Receptors and TREM-1 signalling (Figure 3C) and down regulated pathways associated with cellular motility; actin-based motility of Rho, actin cytoskeletal signalling and Cdc42 signalling. Pathway enrichment analysis identified OxPhos, cholesterol biosynthesis and glycolysis to be among the top-ranked pathways in 6 h PEC monocytes vs 2 h PEC monocytes (Figure 3D). Pseudo-bulk sequencing revealed numerous pathways to be enhanced between each of the experimental groups, OxPhos was upregulated in 2 h blood monocytes compared to naïve blood, and 6 h PEC monocytes compared to 2 h PEC monocytes. Our results demonstrate that OxPhos, could be important in migration (monocytes) and differentiation (monocyte to macrophage). Heat map analysis of the top 30 differentially expressed genes from each pair wise comparison naïve blood vs. 2 h blood (Figure 3E), 2h blood vs. 2 h PEC (Figure 3F) and 2 h PEC vs. 6 h PEC (Figure 3G) revealed distinctive patterns of differential gene expression whereby not all the individual cells from each experimental group followed the same expression pattern. Our single cell transcriptional profiling strongly suggests significantly more complexity within the system than could be appreciated by looking at bulk differential expression between experimental groups.

### Early changes in single cell transcriptome predict heterogeneity and divergence in Ly6C^hi^ monocyte differentiation

In order to provide a more refined bioinformatic analysis of our CD11b^+^Ly6C^hi^ monocyte populations we used Monocle 3 which can predict pseudo-time cell trajectories based on a single timing input of the root cells population (i.e. naïve blood monocytes). An independent clustering algorithm within Monocle 3 software confirmed our prior clustering of cells into 9 discrete clusters over our 4 experimental groups (Figure 4A/B). The initial prediction suggests two discrete cell trajectories. This analysis segregated cells based on the experimental group the cells were from e.g. naïve blood or the *in vivo* compartment (blood vs peritoneum) (Figure 4D). Importantly, Monocle 3 did not have access to other cells group information (i.e. the time of harvest or location of where the cells where from).

**Figure 4:**
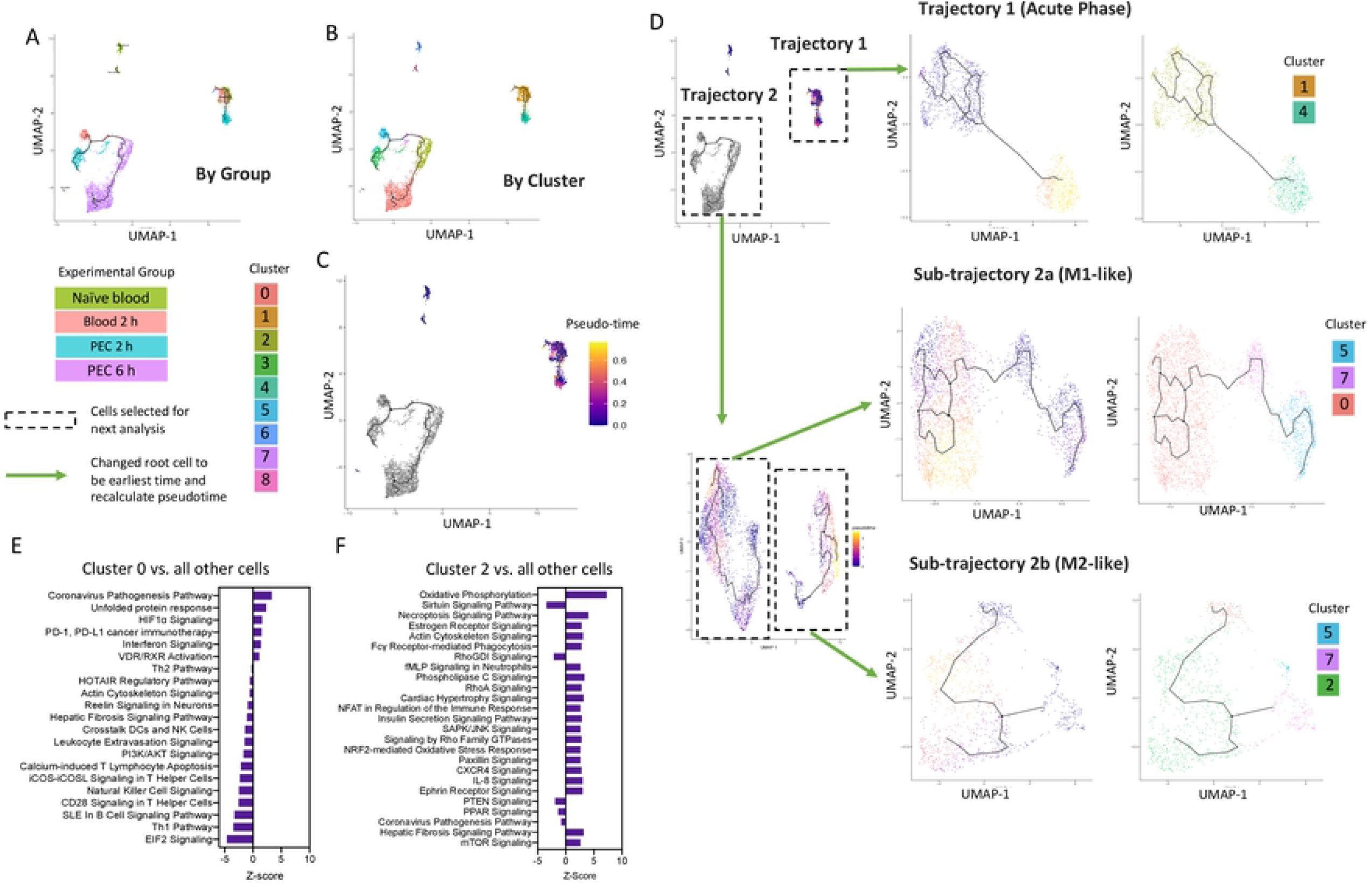
UMAP visualisation of Ly6C^hi^ monocyte differentiation trajectories in the first 6 hours following zymosan challenge. UMAP visualization annotation based on groups **(A)** and clusters **(B)** and by pseudotime **(C) (D)** Iteratively we reanalysed each of the two major trajectories shown in **(B)**. Reanalysis of Trajectory 2 revealed two sub-trajectories which were each re-analysed (lower right). Results of re-analyses are shown in pseudo-time and by cluster. **(E)** Pathway enrichment analysis of cells in Cluster 0 vs. all other cells. **(F)** Pathway enrichment analysis of cells in Cluster 2 vs. all other cells.

The first analysis demonstrated that there was one trajectory (named Trajectory 1) from cluster 1 present in naïve blood, and 2 h blood samples, which projects with a strong pseudo-time prediction score to cluster 4, which is present exclusively in 2 h PEC cells. (Figure 4E). Strikingly, these cells express high levels of IL-7R, and do not follow any further differentiation beyond 2 h; cluster 4 cells are also not present within the peritoneum by the 6 h time point.

Sub-trajectory 2a was a linear progression of cells from cluster 5 present in the 2 h blood progress to cells present in cluster 3 in the 2 h PEC group and ended in cluster 0 present in the 6 h PEC group (Figure 4E). Pathway enrichment analysis using IPA software revealed that cells in Cluster 0 have enrichment for genes related to coronavirus pathogenesis, HIFα and interferon signalling (Figure 4F). Network analysis was performed of the top 250 differentially expressed genes from Cluster 0 vs all other cells, confirmed key involvement of genes known to be highly expressed in M1 macrophages including key transcription factors STAT4 and IRF7 (Sup Figure 4A/B). Suggesting that these are a pro-inflammatory subset of Ly6C^hi^ monocytes. Sub-trajectory 2b is a direct progression of cells from cluster 7 present in the 2 h blood to become cells in cluster 2 present in the 6h PEC monocytes following zymosan challenge (Figure 4E). Pathway enrichment analysis predicted cells in cluster 2 have positive signatures for pathways such as OxPhos, fMLP signalling and NFAT regulation of immune responses (Figure 4G). Network analysis was performed of the top 250 differentially expressed genes from Cluster 2 vs all other cells confirmed key involvement of genes known to be highly expressed in M2 macrophages including Arg1 and IL4R (Sup Figure 4C/D). These pathways are known to be important for M2 macrophages differentiation and regulation of myeloid anti-inflammatory functions and inflammation resolution. These analyses tell us for the first time that as early as 2 h after inflammatory stimuli Ly6C^hi^ monocytes are pre-programmed in the blood to have a distinct cellular and functional fate at the site of inflammation.

### Oxidative phosphorylation is required for monocyte/macrophage chemotaxis

The process of monocyte migration requires ATP production and hydrolysis for actin polymerisation via Rac/RhoA/Cdc42, cell membrane re-arrangement and biosynthesis which are essential for directional movement. The top enriched pathways between naïve blood and 2 h Blood monocytes was OxPhos, suggesting that OxPhos is the mechanism by which naïve blood monocytes generate the ATP needed for migration to site of inflammation. To test this hypothesis, we utilsed an *ex vivo* real time chemotaxis system to investigate the role OxPhos in monocyte/macrophage chemotaxis.

Firstly, we wanted in inhibit ATP production from OxPhos and determine if this affected monocyte chemotaxis. We have previously shown that BioGel elicited monocyte/macrophages can undergo chemotaxis towards CCL2 and CCL5 (Iqbal et al., 2013) while Bone-marrow derived macrophages (BMDM) undergo chemotaxis towards C5a (Purvis et al., 2021). Oligomycin inhibits the ATPase F0/F1, which block ATP production via the electron transport chain (Figure 5A). Pre-treatment with oligomycin (1 μM; 15 mins prior to chemotaxis) significantly inhibited BioGel elicited monocyte/macrophage chemotaxis towards CCL2 (Figure 5B) and CCL5 (Sup Figure 5A), and BMDM chemotaxis towards C5a (Sup Figure 5B) assessed by Cell Index Max-Min and area under the curve. Similar results were obtained by using inhibitors of complex I, II, III and IV (Figure 5C-F), suggesting that ATP production from intact electron transport chain via OxPhos is critical for monocyte chemotaxis.

**Figure 5:**
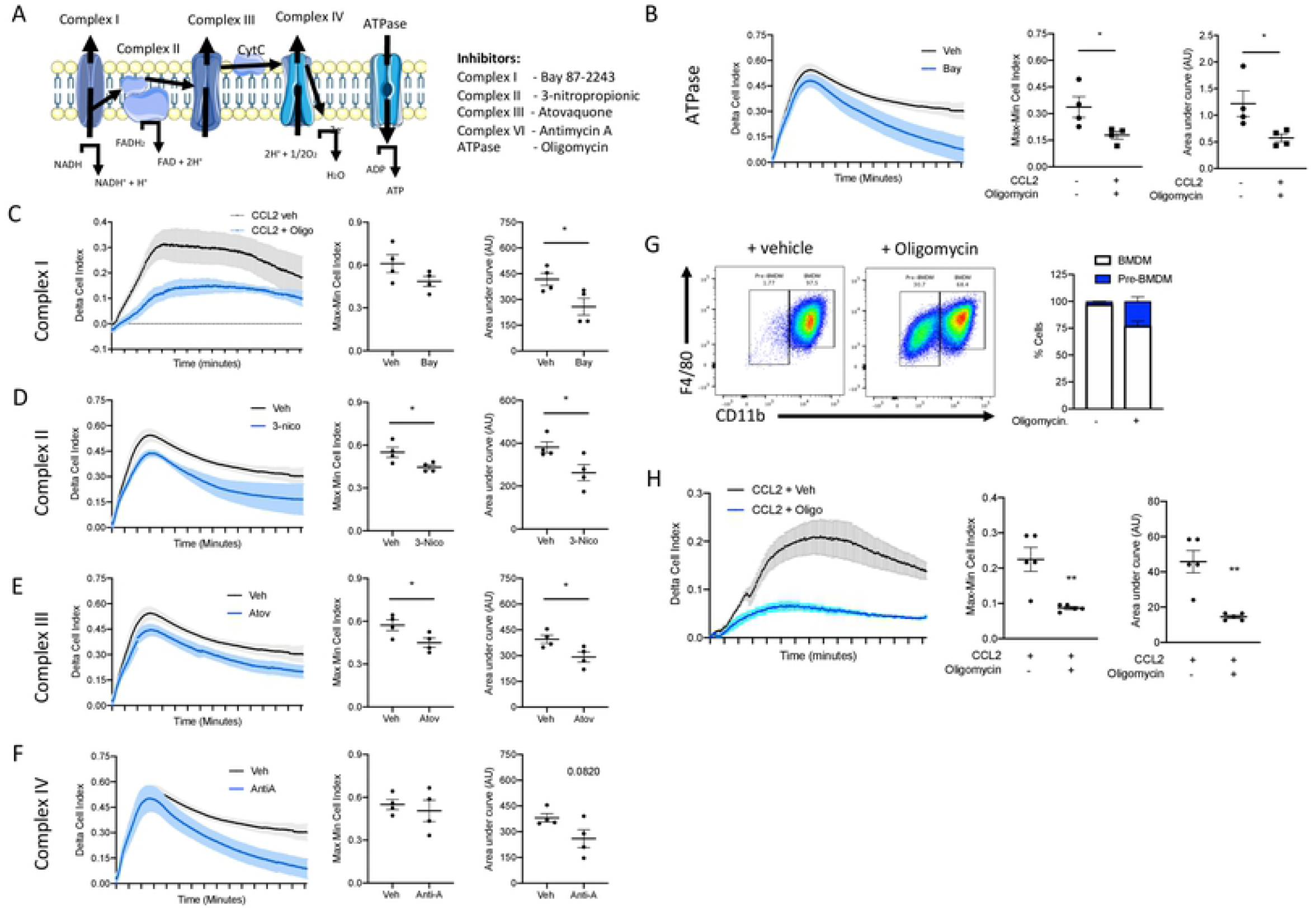
ATP production from Oxidative phosphorylation is essential for monocyte chemotaxis and monocyte to macrophage differentiation. Murine and human monocytes/macrophage need oxidative phosphorylation for monocyte chemotaxis and monocyte to macrophage differentiation **(A)** Schematic of the mammalian electron transport chain. Biogel elicited myeloid cells were incubated with **(B)** oligomycin (1 μM), **(C)** Bay87-2243 (100 nM), **(D)** 3-nitropropionic (1 mM), **(E)** Atovaquine (5 μM) or **(F)** Antimycin A (0.5 μM) and vehicle for 15 min before being added to the upper chamber (4 × 10^5^/well) of a CIM-16 plate and allowed to migrate for 6 hr toward murine CCL2 (10 nM). Cell index was measured at 30 sec intervals for 6 hours and quantified as Max-Min or area under the curve (AUC) analysis. **(G)** Bone marrow monocytes were isolated using immunomagnetic sorting and differentiation into BMDM using L-cell conditioned medium for 7 days. Representative flow cytometry of BMDM treated with oligomycin (1 μM) or vehicle for 15 min before being stimulation with L-cells conditioned medium. Percentage of CD11b^hi^F4/80^+^ and CD11b^lo^F4/80^+^ macrophages was quantified. **(H)** Human monocytes were isolated from buffy coat. 4 × 10^5^/well purified human monocytes were incubated with oligomycin (1 μM) or vehicle for 15 min before being added to the upper chamber of a CIM-16 plate and allowed to migrate for 8 hr toward human CCL2 (10 nM). Migration was measured with max-min analysis and area under the curve (AUC) analysis. Statistical analysis was conducted by one-way ANOVA with Dunnett’s multiple comparison post-test. *p=0.05, **p=0.01, ***p=0.001, ****p=0.0001.

### Oxidative phosphorylation is required for murine monocyte to macrophage differentiation

Having demonstrated that energy metabolism via OxPhos is one of the top ranked canonical pathways to be switched on following inflammatory stimulus we questioned whether OxPhos was needed for monocyte to macrophage differentiation. Monocytes were isolated from mouse bone marrow using magnetic separation and cultured for 7 days in L-929 conditioned media (containing M-CSF) to differentiate into BMDM and cells were analysed by flow cytometry using the cell surface markers CD11b and F4/80. To test if ATP production from OxPhos is needed for monocyte to macrophage differentiation monocytes were treated with 1 μM oligomycin or vehicle during the 7-day differentiation protocol. Treatment with oligomycin significantly reduced the formation CD11b^+^F4/80^+^ macrophages confirming that ATP production from OxPhos is essential for monocyte to macrophage differentiation (Figure 5G).

### Oxidative phosphorylation is required for human monocyte chemotaxis

We next wanted to confirm the results we obtained in primary murine monocytes in primary human monocytes, and human monocyte derived macrophages. Human monocytes were immunomagnetically sorted from peripheral blood mono-nuclear cells isolated from leukocytes cones. Primary human monocytes were pre-treated with the ATPase F0/F1 inhibitor oligomycin (1 μM) for 15 mins prior to being added to the top chamber of the CIM-16 plate. Inhibiting ATP production via OxPhos inhibited primary human monocyte ability to undergo chemotaxis towards CCL2 (Figure 5H), analysed using both the maximal rate of chemotaxis (Max-Min CI) but also the total migration (AUC) (Figure 5H). Collectively, these results are consistent with our single cell RNA Sequencing data that suggested a key role for OxPhos in monocytes migration and monocyte to macrophage differentiation in the early stages of the inflammatory response.

## Discussion

In the present study we sought to better understand at a single cell level the earliest transcriptional events determining blood monocyte fate as they are recruited to, and differentiate within a site of resolving inflammation. Our experiments reveal there is significant heterogeneity within the circulating Ly6C^hi^ monocyte populations, and that Ly6C^hi^ monocytes are pre-programmed in the blood to become either M1-like or M2-like macrophages at a site of inflammation. Critically we have demonstrated that there is a rapid and dynamic metabolic shift toward oxidative metabolism in the sub-set of blood Ly6C^hi^ monocytes that are predicted to become M2-like macrophages. This metabolic shift occurs earlier than has been demonstrated before, specifically prior to displaying classical macrophage markers, which are maintained as they differentiate. Indeed, we demonstrate for the first time a requirement for ATP production from intact electron transport chain for both monocyte/macrophage chemotaxis and monocyte to macrophage differentiation.

Under steady state it was widely accepted that tissue resident macrophage, self-maintain through-out adult life with minimal contribution from circulating monocytes (Yona et al., 2013)(Hashimoto et al., 2013)(Watanabe et al., 2019). However, this dogma has being challenged. Most tissues are now recognized to contain multiple macrophage populations localised to their distinct microanatomical domains (Gibbings et al., 2017)(Tabula Muris Consortium et al., 2018). Each of these myeloid cell populations differs in its ontogeny, capacity for self-renewal, and that each macrophage population likely plays a specialised role in tissue homeostasis, tissue injury, and tissue repair (Rahman et al., 2017)(Dick et al., 2019). Critically their rate of replacement by monocyte-derived cells is also vastly different under steady state. Until now current methods have also had limited success in tracking monocyte progeny in inflamed tissues and it was still unclear whether monocyte-derived macrophages transiently infiltrate inflamed tissues or persist locally once the inflammation resolves (Yona et al., 2013). Our lineage tracing experiments demonstrate that following peritoneal inflammation the new peritoneal macrophages differentiate from blood derived Ly6C^hi^ monocytes and persist in the peritoneum to differentiate into either M1-like or M2-like macrophages.

Unbiased tSNE visualisation based on Louvain clustering of single cell RNA sequencing data revealed there is significant heterogeneity within inflammatory monocytes (CD45^+^CD11b^+^Ly6G^-^Ly6C^hi^) isolated from our 4 experimental conditions. Conventional cell surface expression identifies our input cells as homogenous population therefore, it is striking that we can identify significant heterogeneity within the cells present not only between the four experimental groups but within each experimental group. This underappreciated heterogeneity within CD11b^+^Ly6C^hi^ monocytes defined by flow cytometry gives rise to divergent functions. Our experiments further demonstrate that Ly6C^hi^ monocytes are pre-programmed to become one of two major effector cells in the inflamed peritoneum.

Importantly, our data suggests that differentiation into a monocytes-derived macrophage is pre-programmed prior to receiving an inflammatory stimulus. This finding does not agree with the current convention that monocytes differentiate into macrophages, and then polarise following the activating signal. Our data also collaborates a recent study identified at least two unique tissue-resident interstitial macrophages in the steady-state lung that could be distinguished by unique transcriptional profiles and spatially localized to the interstitium of the bronchovascular bundles, but not alveolar walls. In line with our findings it was identified that other tissue resident macrophages are blood monocyte derived at both steady state and following inflammatory stimulation (Gibbings et al., 2017)(Evren et al., 2021). Others have shown using transcriptome-based network analysis in a spectrum model of human macrophage activation, that macrophages *in vivo* do not fully fall into the classical M1/M2 paradigm (Xue et al., 2014)(Nahrendorf and Swirski, 2016). Our data adds compelling evidence that post inflammatory tissue niches can be repopulated by a diverse range of functionally distinct Ly6C^hi^ monocyte-derived macrophages, and that these may be identified by their metabolic signature.

Our experiments have discovered two main trajectories by which CD11b^+^Ly6C^hi^ monocytes differentiate in the first 6 h after inflammatory stimulus. Trajectory 1 is characterised as containing IL-7R^hi^ monocytes. This trajectory had its origins in cells that are present in the naïve blood, suggesting that these are most likely the first monocytes to be recruited following a sterile inflammatory stimulus. IL-7 is produced by numerous cells types including the thymocytes, hepatocytes and stromal cells. To our knowledge IL-7 levels either local or circulating have not been measured in zymosan induced peritonitis, however pam3csk4 (a wildly used TLR2 agonist) is able to induce IL-7R on CD14^+^ monocytes. More recently it has been reported that IL-7R is a key transcriptional signature in patients with inflammatory bowel disease that is unresponsive to anti-TNFα therapies (Belarif et al., 2019). While others have shown that TNFα regulates IL-7R expression on human monocytes (Al-Mossawi et al., 2019). Our data suggests that IL-7R monocytes may be critical in the very early response to systemic cytokines in models such as sepsis. TNFα is also chronically elevated in diseases such as T2DM/obesity and rheumatoid arthritis, so there could be a central role of IL-7R^+^ monocytes in responding to TNFα signals in chronic diseases and hence IL-7 and its receptor could represent a new therapeutic targets.

Our findings demonstrate that Ly6C^hi^ monocytes rapidly up-regulate key genes in the OxPhos pathway, within the first 2 h in blood monocytes. This suggested a need for more efficient and increased ATP production during the acute activation phase in the blood prior to mobilisation to the site of inflammation. We therefore sought to determine if ATP production from OxPhos is needed for monocyte chemotaxis to a range of physiological monocytes/macrophage chemokines using a real time chemotaxis platform (Rumianek and Greaves, 2020)(Iqbal et al., 2013). Indeed, we were able to demonstrate that when the ATPase F0/F1 was inhibited with oligomycin, this reduces the bioavailability of ATP and there was significantly reduced murine monocyte chemotaxis to towards CCL2/CCL5; and a similar result was obtained in human monocytes. The remodelling of the cytoskeleton to enable cell motility consists of actin polymerisation and actomyosin contraction that is sustained through ATP and GTP hydrolysis (Parsons et al., 2010). We, therefore, extended our investigation to determine if disruption of the electron transport chain at different points could also alter monocytes chemotaxis, and found that disruption of complex I, II, III and IV using chemical inhibitors also resulted in reduced ability of monocytes to undergo chemotaxis. These data are the first to suggest a requirement of ATP production from an intact electron transport chain/OxPhos for monocyte chemotaxis.

Our datasets further revealed a transitional monocyte stage with a gene signature indicating active protein synthesis and turnover. The transition from quiescence (naïve blood) to cellular differentiation (2 h PEC) is dependent on increased ribosome biogenesis and protein synthesis (Sanchez et al., 2016) before ribosomal genes are suppressed during terminal differentiation (6 h PEC) (Athanasiadis et al., 2017). We confirmed this experimentally, showing that a mixed bone marrow population which included hemopoietic stems cells and monocytes, failed to fully differentiate into macrophages (CD11b^+^F4/80^+^) in the presence of oligomycin, reinforcing the idea that there is a critical need for OxPhos for energy dependant processes like protein synthesis. Pathway enrichment analysis of pseudo-bulk sequencing between experimental groups revealed a central role for OxPhos at two critical time points 2 h blood monocyte and naïve monocytes, and between 2 h Peri Mono and 6 h Peri Mono. This is important and but also when taken in context with our cluster analysis we see that globally OxPhos is not up-regulated in all monocyte clusters in each of the experimental groups, but rather is restricted to monocytes present in sub-trajectory 2b (Figure 3D). Only by using a single cells RNA-transcriptomics can the subtilties and inherent heterogeneity of the mononuclear phagocyte system be fully appreciated.

During the resolution phase of atherosclerotic plaque regression in animal models continued recruitment of Ly6C^hi^ monocytes into the lesion is essential for plaque regression. These recruited Ly6C^hi^ monocytes then undergo M2-like conversion in a STAT6 dependent manner (Rahman et al., 2017). However, in the setting of hyperlipidaemia monocytes have decreased ability to undergo efficient OxPhos (Baardman et al., 2018). Our data demonstrate for the first time OxPhos is essential for a) Ly6C^hi^ monocytes recruitment to the site of inflammation and b) OxPhos is again needed for M2-like polarisation. This could be why in non-resolving atherosclerotic lesions there is reduced Ly6C^hi^ monocyte recruitment, which is essential for plaque regression and increased M2-like polarisation.

In conclusion we have demonstrated that there is significant previously unrecognised heterogeneity in Ly6C^hi^ monocytes. Single cell transcriptomics of Ly6C^hi^ monocytes coupled with pseudo-time trajectory analysis revealed two distinct differentiation trajectories by which Ly6C^hi^ monocytes differentiate into monocyte-derived macrophages following inflammatory stimuli. We provide the first evidence that monocytes are pre-programmed in the blood to become distinct and functionally diverse macrophage populations. We have also revealed that there is a requirement for a rapid metabolic reprogramming in a subset of Ly6C^hi^ monocytes in the blood within 2 h, which allows them to commit along a differentiation trajectory towards an M2-like phenotype. Opening up the possibility to have targeted prophylactic-therapies that alter monocyte metabolic capacity to facilitate enhanced macrophage M2-like polarisation to aid inflammation resolution and tissue repair if needed.

## Acknowledgements

This work was funded by British Heart Foundation (BHF) Programme Grant awards to DRG and KMC (RG/15/10/31485 and RG/17/10/32859) and a BHF Chair Award (CH/16/1/32013) to KMC. The transcriptomic analysis was supported by the Wellcome Trust Core Award Grant Number 203141/Z/16/Z with additional support from the National Institute for Health Research (NIHR) Oxford Biomedical Research Centre. The views expressed are those of the author(s) and not necessarily those of the NHS, the NIHR or the Department of Health. We thank the Oxford Genomics Centre at the Wellcome Centre for Human Genetics (funded by Wellcome Trust grant reference (203141/Z/16/Z) for the generation and initial processing of the sequencing data.

Conceptualisation, GP, EM, KC,DRG; Methodology, GP and EM; Formal Analysis: GP, EM, BW, SR; Investigation, GP and EM; Writing First draft, GP, EM, KC, DRG; Writing Review and Editing, GP, EM, KC, DRG, BW; Visualisation, GP, BW; Supervision, EM, DRG, KC, HL; Funding Acquisition, KC and DRG.

---

| REAGENT or RESOURCE | SOURCE | IDENTIFIER |
| --- | --- | --- |
| **Experimental Models: Organisms/Strains** |  |  |
| C57BL/6 | Charles River | C57BL/6NCrl |
| L929 cells | ATCC | ATCC NCTC clone 929 |
| Human peripheral blood mononuclear cells | NHSBTS | n/a |
| hCD68-GFP | Generated in-house | n/a |
| **Antibodies** |  |  |
| BV421-conjugated CD11b (anti-mouse) | BioLegend | 101236 |
| BV510-conjugated Ly6G (anti-mouse) | BioLegend | 127633 |
| PerCP-conjugated Ly6C (anti-mouse) | BioLegend | 128028 |
| APC-conjugated F4/80 (anti-mouse) | BioLegend | 123116 |
| PE/Cy7-conjugated CD115 (anti-mouse) | BioLegend | 135524 |
| Fc-blocking anti-CD16/32 (anti-mouse) | BioLegend | 101302 |
| PE/Cy7-conjugated CD71 (anti-mouse) | BioLegend | 113812 |
| FITC-conjugated CD206 (anti-mouse) | BioLegend | 141704 |
| PE-conjugated MHCII (anti-mouse) | BioLegend | 107608 |
| PE-conjugated CD16 (anti-human) | BioLegend | 360704 |
| FITC-conjugated CD14 (anti-human) | BioLegend | 325604 |
| BV421-conjugated CD11b (anti-human) | BioLegend | 101236 |
| APC-conjugated CD71 (anti-human) | BioLegend | 334108 |
| FITC-conjugated CD206 (anti-human) | BioLegend | 321104 |
| PerCP-conjugated HLA-DR (anti-human) | BioLegend | 307628 |
| BioLegend Cell Hashing A | BioLegend | 155861 |
| **Chemicals, Peptides, and Recombinant Proteins** |  |  |
| Recombinant murine Interferon $\gamma$ | PeproTech | Cat# 315-05 |
| Lipopolysaccharide | Sigma | Cat# L4391 |
| Recombinant human mCSF | peproTech | Cat# 300-25 |
| Recombinant murine IL-4 | peproTech | 214-14 |
| Recombinant human IL-4 | peproTech | 200-04 |
| Recombinant Human Interferon $\gamma$ | PeproTech | 300-02 |
| oligomycin | Sigma/Merck | 75351-5MG |
| FCCP | Sigma/Merck | C2920-10MG |
| Antimycin A | Sigma/Merck | A8674-50MG |
| Rotenone | Sigma/Merck | R8875-1G |
| Recombinant murine C5a | peproTech | 315-40 |
| Recombinant murine CCL2 | peproTech | 250-10 |
| Recombinant murine CCL5 | peproTech | 250-07 |
| Recombinant human CCL5 | peproTech | 300-06 |
| **Oligonucleotides** |  |  |
| *Arg1* | Qiagen | QT00134288 |
| Rel | Qiagen | QT00245359 |
| nos2 | Qiagen | QT001547980 |
| 18s | Qiagen | QT02448075 |
| B-actin | Qiagen | QT00095242 |
| **Critical Commercial Assays** |  |  |
| Huma Monocytes isolation kit | Miltenyi Biotec | 5170524422 |
| Murine bone marrow monocyte isolation kit | Miltenyi Biotec | 52003003994 |
| Chromium Single Cell 3' Reagents Kits | 10X Genomics | PN-120237 |
| Qubit dsDNA HS Assay Kit | Thermofisher | Q32851 |
| Agilent High Sensitivity D5000 ScreenTape | Agilent | 5067- 5592 |
| D1000 ScreenTape | 5067- 5582) | 5067- 5582 |
| QuantiTect Reverse Transcription kit | Qiagen | 205313 |
| Sybr Select gene expression master mix | Applied Biosystems | 4309155 |
| **Specialist Machines** |  |  |
| autoMACS Pro Separator | Miltenyi Biotec | n/a |
| Seahorse XFe96 analyser | Agilent | n/a |
| Chromium 10x Genomics platform | 10X Genomics | n/a |
| Illumina HiSeq 4000 | Illumina | n/a |
| xCELLigence RTCA-DP | Agilent | n/a |
| Nanodrop | Thermo Scientific | n/a |
| StepOnePlus™ thermal cycler | Applied Biosystems | n/a |
| BD Fortessa X20 cytometer | BD Biosciences | n/a |
| DAKO CyAn cytometer | Beckman | n/a |
| **Software and Algorithms** |  |  |
| Graph Pad Prism 7 | Graph Pad | https://www.graphpad.com/scientific-software/prism/ |
| FlowJo | Tree Star Inc |  |
| Diva software | BD Biosciences |  |
| Wave (Seahorse) | Agilent |  |

**Supplementary Figure 1:**
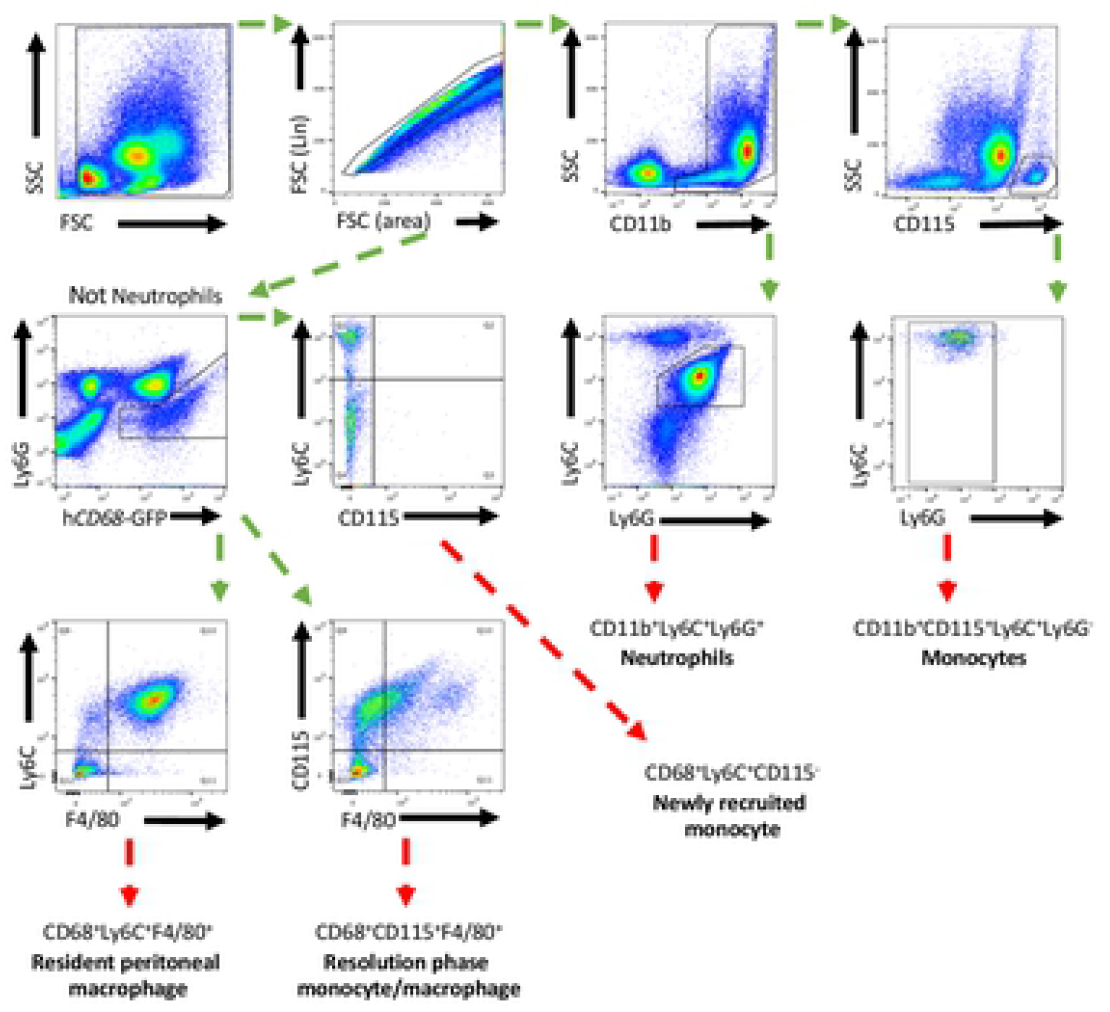
Full gating strategies for myeloid cells present in the peritoneum.

**Supplementary Figure 2:**
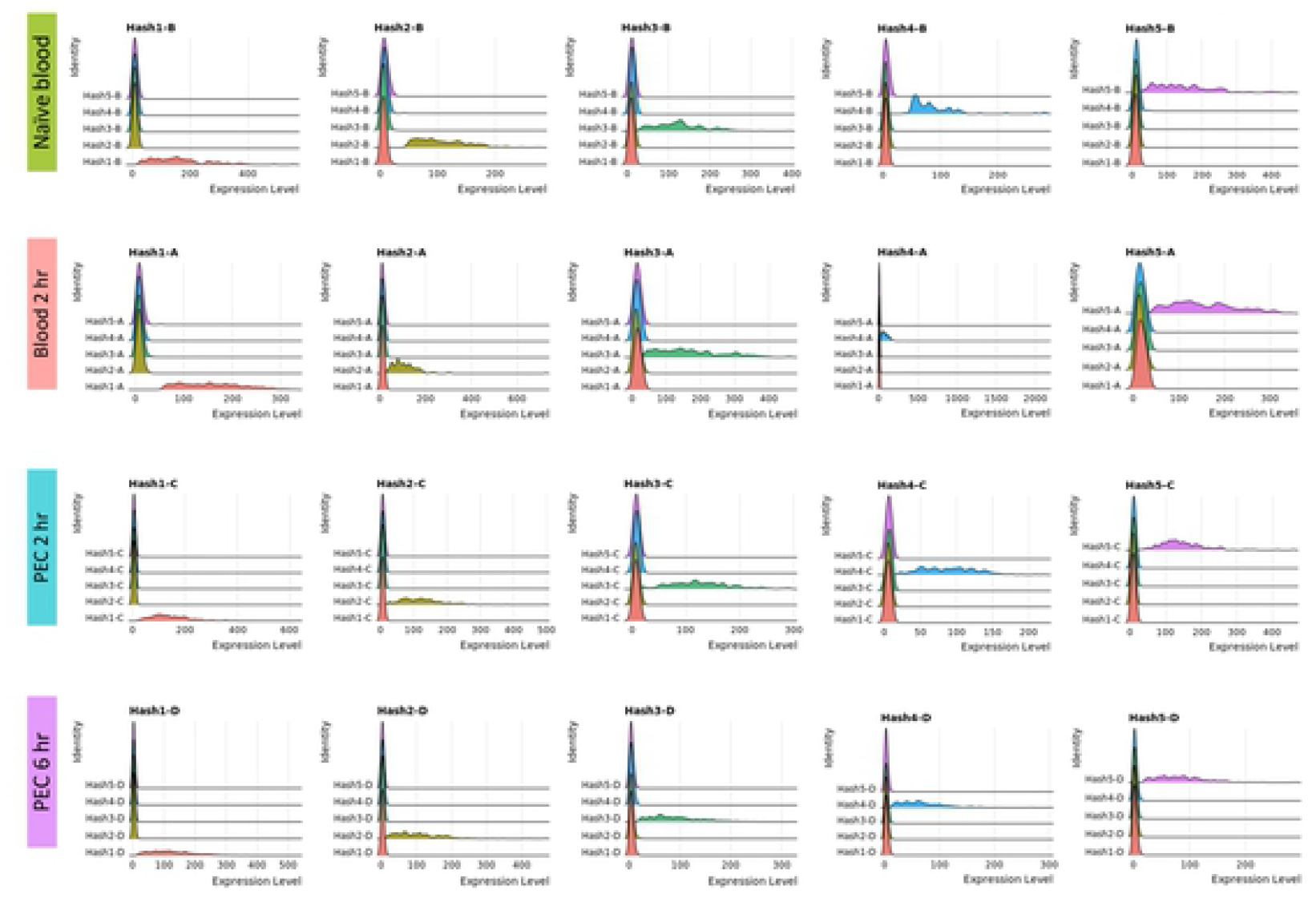
Ridge plot analysis for hashing antibody efficiency. Initial quality control of cell hashing and clustering algorithm.

**Supplementary Figure 3:**
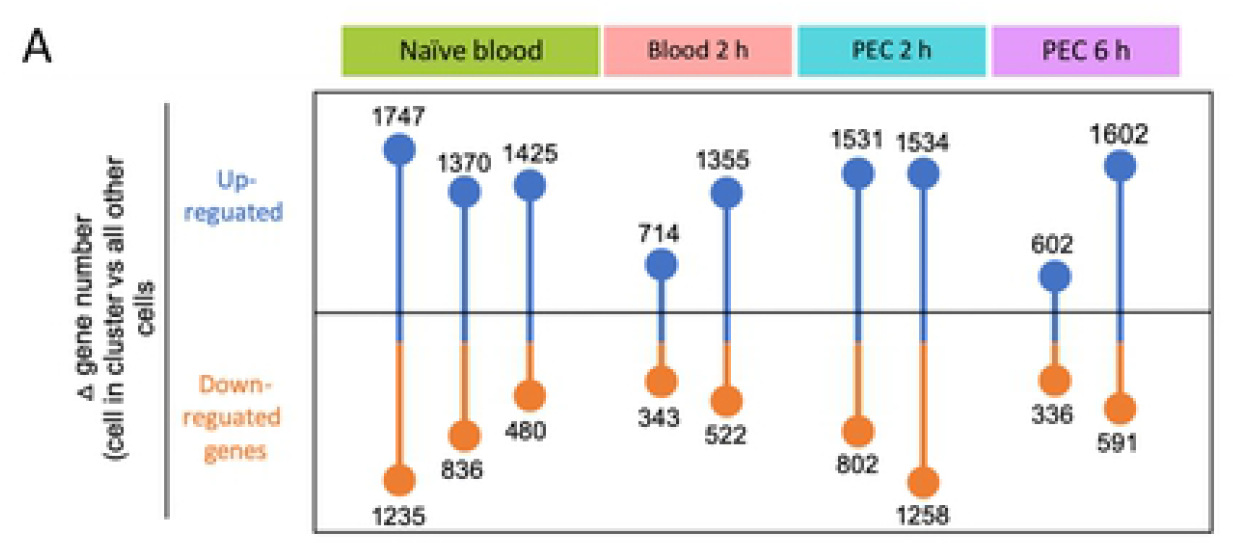
Single cell RNA sequencing reveals significant differential gene expression between Ly6C^hi^ monocyte clusters.

**Supplementary Figure 4:**
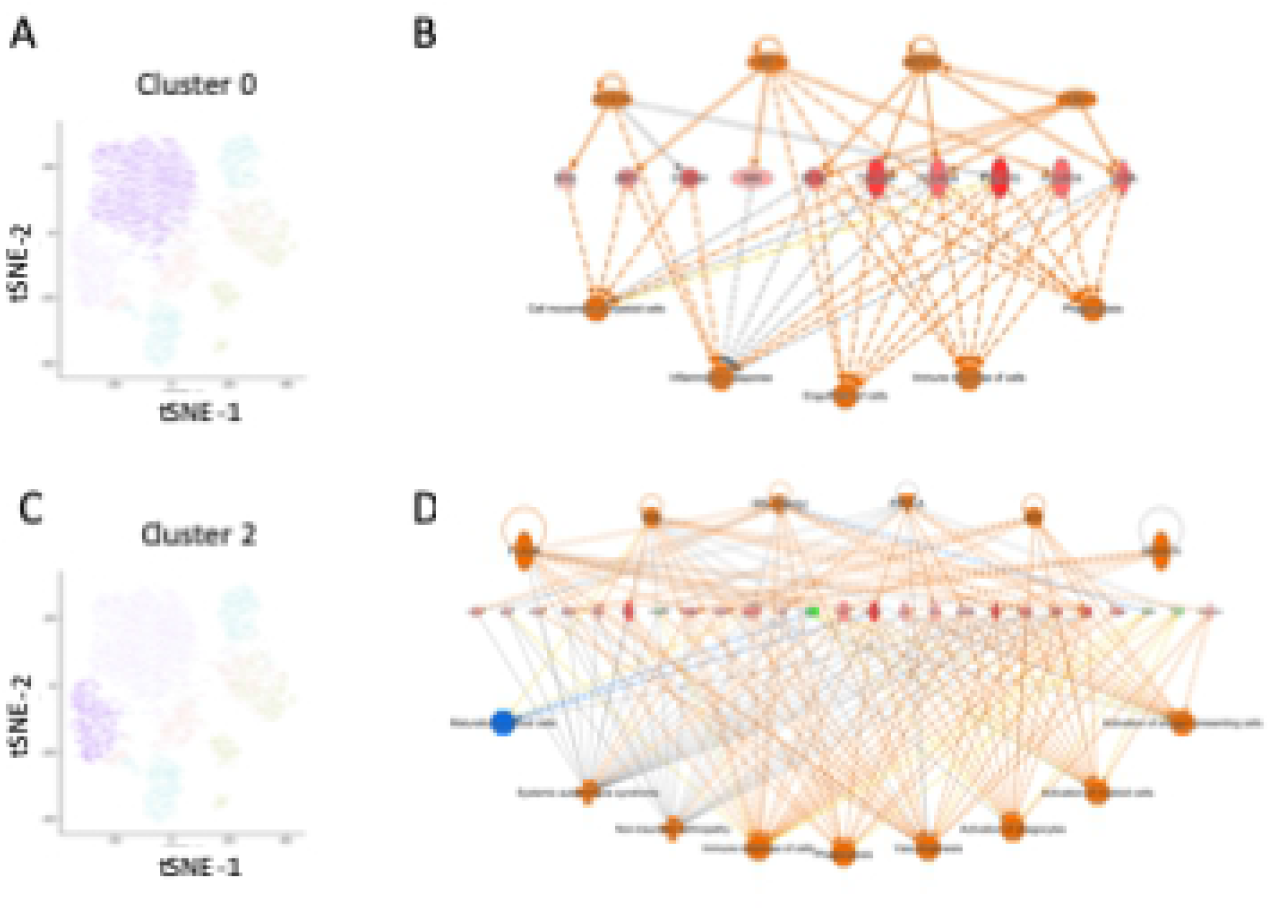
Transcript network analysis reveals monocytes are programmed to have a pro-inflammatory or resolving phenotype as early as 6 h following zymosan challenge.

**Supplementary Figure 5:**
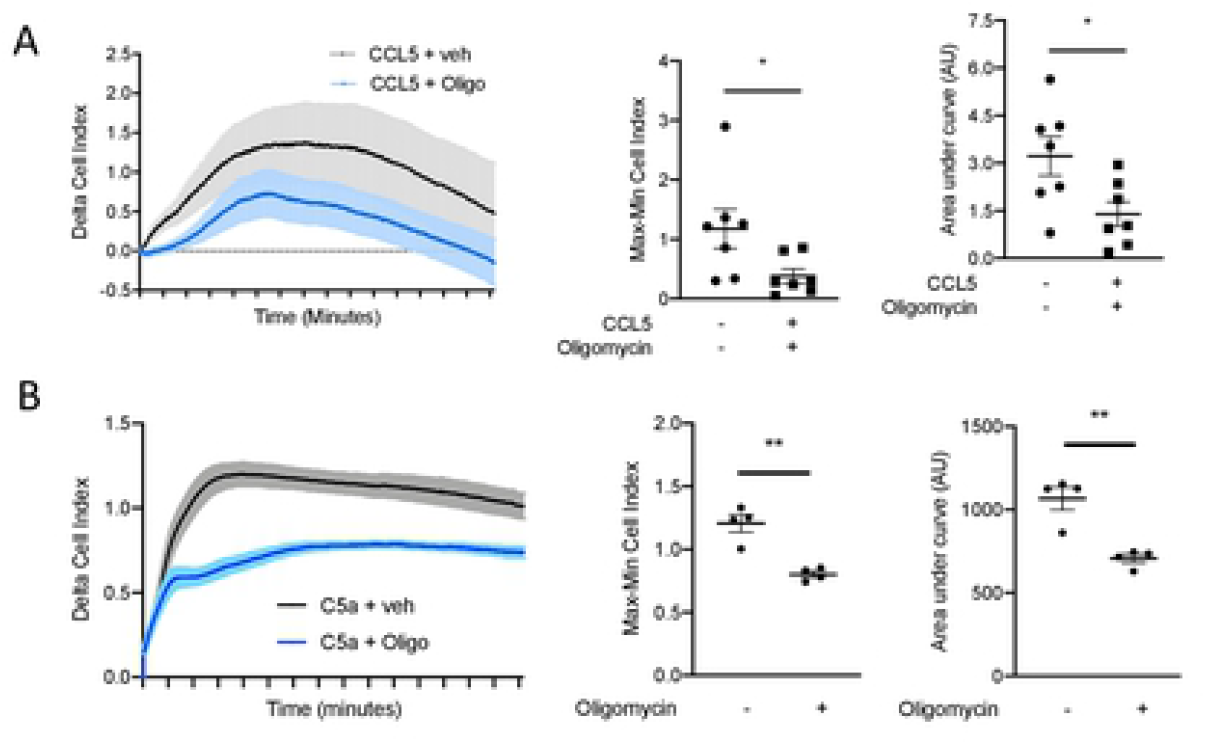
ATP production from oxidative phosphorylation is essential for monocyte/macrophage chemotaxis. **(A)** Biogel elicited myeloid cells were incubated with oligomycin (1 μM or vehicle for 15 min before being added to the upper chamber (4 × 10^5^/well) of a CIM-16 plate and allowed to migrate for 6 hr toward murine CCL5 (10 nM). Cell index was measured at 30 sec intervals for 6 hours and quantified as Max-Min or area under the curve (AUC) analysis. **(B)** Bone marrow derived macrophage were incubated with oligomycin (1 μM or vehicle for 15 min before being added to the upper chamber (4 × 10^5^/well) of a CIM-16 plate and allowed to migrate for 6 hr toward murine complement C5a (10 nM). Cell index was measured at 30 sec intervals for 6 hours and quantified as Max-Min or area under the curve (AUC) analysis. Statistical analysis was conducted by one-way ANOVA with Dunnett’s multiple comparison post-test. *p=0.05, **p=0.01.

